# GIA: A genome interval arithmetic toolkit for high performance interval set operations

**DOI:** 10.1101/2023.09.20.558707

**Authors:** Noam Teyssier, Martin Kampmann, Hani Goodarzi

## Abstract

**Motivation:** This study addresses the pressing need for efficient interval techniques in processing vast genomic datasets, such as those generated by ChIP-seq, RNA-seq, and whole-genome sequencing. Intervals are critical in characterizing biological features, necessitating streamlined interval manipulation for meaningful insights. Existing tools often struggle with memory and runtime requirements when managing extensive genomic region arithmetic.

**Results:** The study introduces GIA (Genomic Interval Arithmetic) and BEDRS, a novel command-line tool and a rust library that significantly enhance the performance of genomic interval analysis. GIA outperforms existing tools like BEDOPS, BEDTools, and GenomicRanges by a factor of 2x to 20x across a range of operations. These advances enable researchers to perform genomic interval operations more efficiently, drastically reducing computational time and resource requirements in high-throughput genomic sequencing analysis.

**Availability and Implementation:**

- https://github.com/noamteyssier/gia
- https://github.com/noamteyssier/bedrs

## Introduction

Genomic data analysis plays a fundamental role in deciphering biological complexities from datasets generated using techniques like ChIP-seq, RNA-seq, and whole-genome sequencing. However, as datasets grow in size, efficiently managing and analyzing genomic regions becomes challenging. While tools like ***BEDTools***(Quinlan, 2014; Quinlan and Hall, 2010), ***BEDOPS***(Neph *et al*., 2012), and ***GenomicRanges***(Lawrence *et al*., 2013) are instrumental and highly optimized, they still face difficulties when handling large datasets, often due to memory usage and processing speed limitations. Additionally, ***BEDTools*** and ***BEDOPS*** are primarily command-line interfaces and their core functionality is not modular and usable by new tools directly.

To address these challenges and drawbacks, we introduce ***gia*** (Genomic Interval Arithmetic), a practical and optimized tool for genomic interval analysis and ***bedrs***, a modular and fully generic genomic interval library written in rust. Our tools maintain the core functions of both ***BEDTools*** and ***BEDOPS*** and can be run in either an in-place memory-hungry approach or in a constant memory streamable manner. Both the command-line tool, ***gia***, and the genomic interval library, ***bedrs***, are presented as free and open-source tools.

## Methods

***gia*** is a command-line tool built in the rust programming language and manages the file IO and library function calls. All the interval operations and logic are performed within the ***bedrs*** interval library.

### bedrs

***bedrs*** is a genomic coordinate interval library built fully generically around the idea of genomic intervals and does not specify a rigid structure for what a genomic interval can be or what types it is composed of. All interval types must implement the *Coordinate trait*, which describes a minimal set of descriptors of what the abstract coordinate structure type is. Traits are a set of functions, described by their outputs, which can be used to define the behavior of an abstract type. Once a new interval structure implements the *Coordinate trait*, it can use the full set of interval operations within the library at no extra cost.

This allows for a highly flexible framework for performing genome interval arithmetic without being limited to a fixed structure, like the BED(Kent *et al*.) format or its variants BED6, BED12, etc. It also allows for easy generalization to BAM(Li *et al*., 2009), GFF(Eilbeck *et al*., 2005), GTF, VCF(Danecek *et al*., 2011), and any other interval-like structure that can implement the basic functions of the *Coordinate* trait. This generalization also fixes performance to the type structure, giving the developer flexibility to tune their performance optimizations by type and thereby allowing more precise optimization by use-case without affecting the performance of other structures. I.e. one can create an interval type with additional fields beyond the minimal set, and use all of the functionality of the library, but never affect the performance of those base intervals operations or the performance of other types within the library without the additional fields.

While ***bedrs*** methods are implemented generically, we do provide built-in interval structures such as a base interval (start, end), a genomic interval (chr, start, end), and a stranded genomic interval (chr, start, end, strand). These built-ins can be thought of as structures with “batteries-included”, and in fact are the structures used within ***gia***. They are highly documented and useful resources for the design of novel standardized interval structures in bioinformatics.

The design choice of a generic trait based genomic interval library is to take advantage of rust’s strong type system and powerful compiler. Each definition of a new interval type implementing the *Coordinate* trait has a monomorphized and optimized set of operations within the compiled binary. Monomorphization is a compilation process to take generic implementations and write optimized machine code for each unique combination of generic types used in the code. This is done to eliminate the overhead of runtime polymorphism and achieve better performance.

These monomorphized implementation improvements come at very little cost (slightly longer compilation time), and in fact come with many benefits, as it allows for a very flexible structure design during algorithm development without sacrificing computational efficiency.

### gia

#### Numerical Representations of Chromosomes

***gia*** takes advantage of the strong typing system in rust to efficiently serialize and deserialize BED files into internal genomic interval types which can be used for set operations. An efficient design choice of ***gia*** is that it can be used either in a *named* or *unnamed* context. This allows for more optimized interval operations when chromosome names are numerical (unnamed) as opposed to string representations (named). Internally, both forms are treated as numeric, but during *named* serialization, ***gia*** calculates and stores a thin mapping of chromosome names to numeric indices - drastically reducing memory requirements and runtimes in most genomic interval contexts. This is because the number of chromosome names is generally orders of magnitude less than the number of intervals, and so keeping a full-sized string buffer for each record is memory-expensive and inefficient. Additionally, comparisons of shared chromosomes with integer representations is an O(1) operation, but comparison of string representations is O(N), where N is the length of the chromosome strings. Precomputing numerical indices drastically reduces memory usage within a run but also drastically reduces comparison times for each interval pair.

#### In-place containers or streaming

One of the advantages of ***gia*** over existing genomic interval libraries is the flexibility it allows in its tools. ***BEDTools*** performs most of its operations by first loading the entire interval library into memory, performing the operation, and then writing the library out afterwards. This is computationally expensive but does not necessarily require a fixed ordinal format for the intervals beforehand (such as merged intervals or sorted intervals). ***BEDOPS*** however is built fully around streaming and most operations expect the intervals to be presorted. It gets around the large memory and computational requirements by keeping intervals on the stack only as long as they are needed for the operation to continue - by keeping track of the ordinal nature of intervals within interval sets. This keeps the memory usage of the operation flat and the number of intervals in the set has no impact on the peak-memory usage of the operation. However, in low set numbers, the run-time and memory requirements of storing the interval set is lower than the run-time and memory requirements of managing the interval streams.

***gia*** has been built to be flexible and allows for both in-place and streaming methods to be used. If the intervals are presorted then the operations can be run using a streaming method, which keeps track of the interval streams and manages interval queues to keep them only for the required amount of time. However, if the intervals are not sorted or more advanced operations and subcommands are required, then the operations can be run by first loading the interval set into memory. For low to mid-sized interval datasets both methods have increased performance than ***BEDTools*** and ***BEDOPS***, but at very high interval datasets, the streaming methods have a much better performance - but ***gia*** still outperforms ***BEDOPS*** in both memory usage and time elapsed.

## Results

All benchmarking was performed on a Macbook Pro M2 Max. Run-time estimates were calculated using ***hyperfine***(Peter, 2023) for command-line tools, ***microbenchmark*** for R tools, and peak-memory estimates were calculated using GNU ***time***.

### General Performance

Globally, ***gia*** outperforms ***BEDOPS, BEDTools***, and ***GenomicRanges*** in (Figure 1). On average all commands ran nearly twice as fast as the existing fastest tool. Where applicable, both the numeric (unnamed) and string (named) versions of ***gia*** were run. The named version incurs a slight runtime increase but is still in the same order of magnitude of the unnamed version and significantly below the runtimes of ***BEDTools*** and ***BEDOPS***.

**Figure 1.**
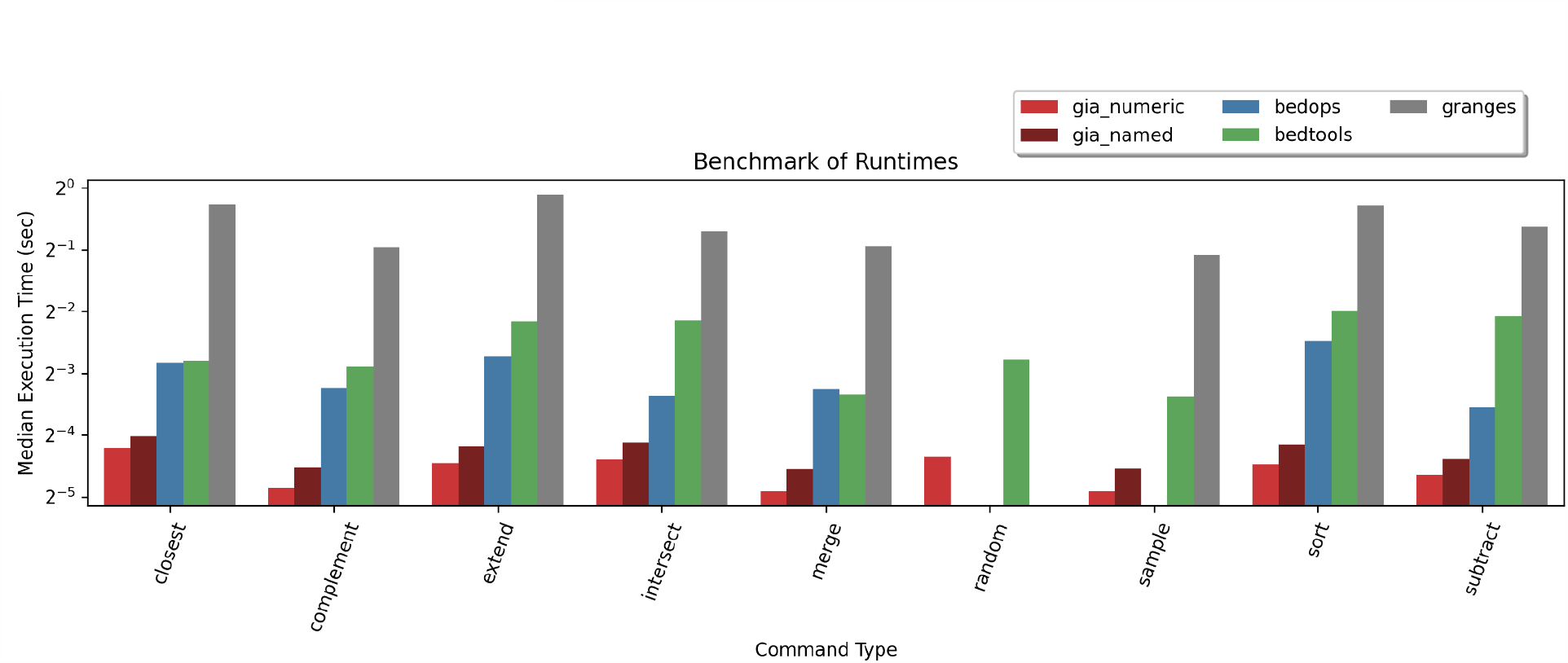
Runtime performance of ***gia, BEDOPS, BEDTools***, and ***GenomicRanges*** (abbreviated ***granges***) over a variety of shared operations between the tools. All operations are performed ‘in-place’ for ***gia*** and **BEDTools** but are performed as streamable operations for ***BEDOPS***. Benchmarks performed on an interval set of size 200,000 sampled randomly across 10 chromosomes. Elapsed time calculated using a Macbook Pro M2 Max.

The performance increases can better be seen by showing the fold-changes compared to the slowest implementation of the operation. For operations that are commonly performed, like subtract, sort, and intersect, we observe that ***gia*** performs 5.5x, 5.2x, and 4x faster than ***BEDTools*** (Figure 2). Compared to ***BEDOPS***, intersect and subtract are still nearly twice as fast, even though the performance is being measured with in-place operations which include the performance cost of a sort as well as the overhead of reading the inputs fully before beginning the operation. Compared to ***GenomicRanges, gia*** is performing at near 15x faster run-times across all operations.

**Figure 2.**
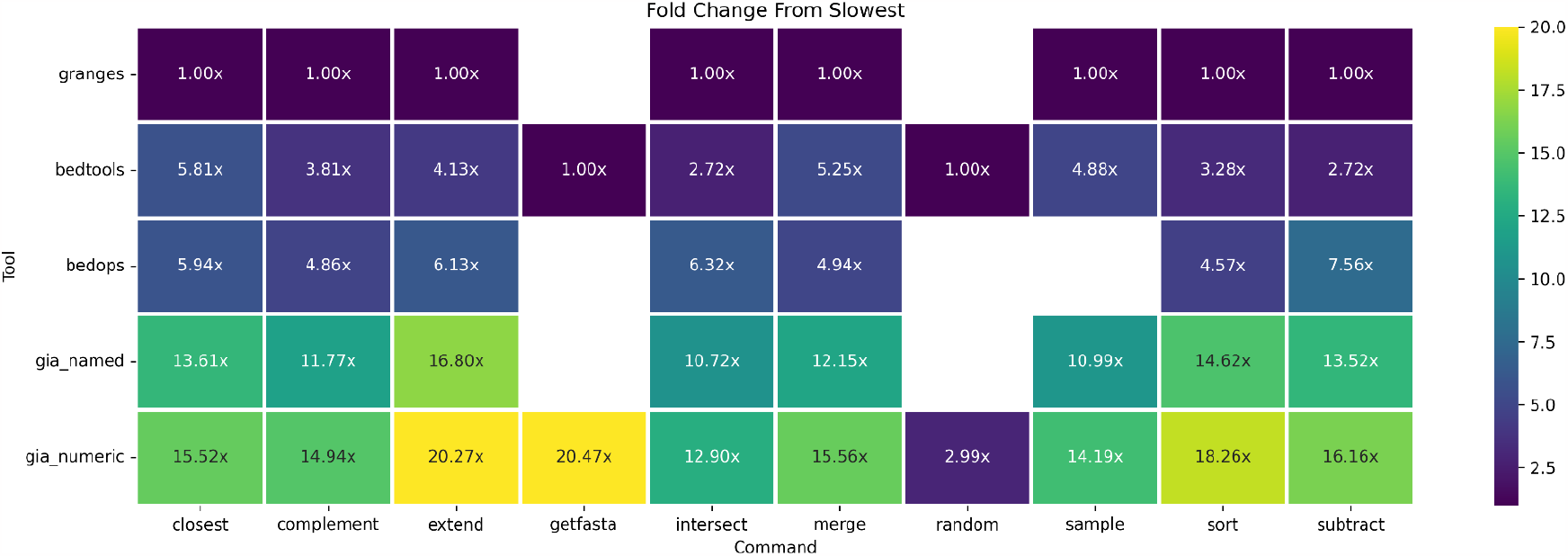
Elapsed time speedup fold-change of each tool compared to the slowest elapsed time for each subcommand. Benchmarks performed on an interval set of size 200,000 sampled randomly across 10 chromosomes.

### Performance Optimization Case Studies

#### get-fasta

A common operation performed with genomic intervals is to extract the genomic sequence defined by some query intervals in a BED file. This operation has been highly optimized within ***gia*** by taking advantage of powerful copy procedures within rust. The fasta file is memory-mapped directly and intervals are used to query the memory map. During interval iteration the genomic sequence can be copied with an O(1) operation without incurring large re-allocation costs for each interval. This results in 20x performance increases compared to the ***BEDTools*** equivalent (Figure 3).

**Figure 3.**
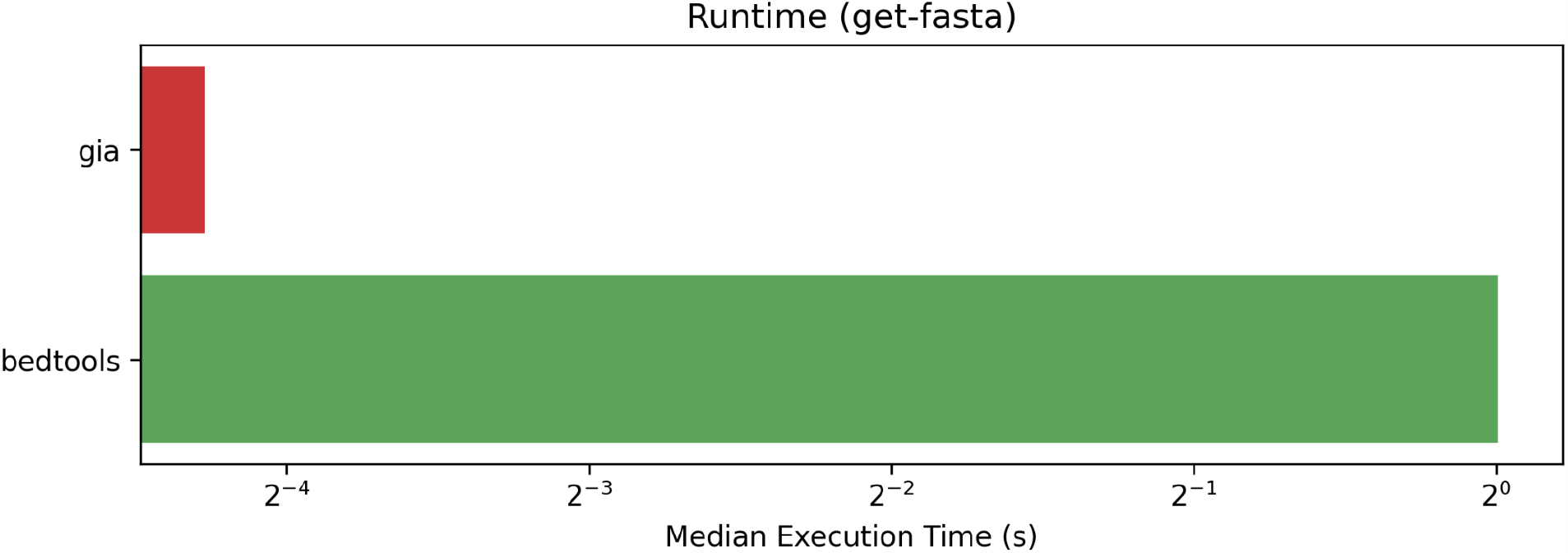
Visualization of elapsed time for running ‘gia get-fasta’ and ‘bedtools getfasta’. Fasta queries were performed on the Ensembl Homo sapiens primary assembly DNA using 200,000 randomly sampled intervals.

#### splici

Another commonly used genomic interval operation is the splici algorithm. This algorithm is utilized by the ***pyroe***(He *et al*., 2022) package in python, and a variant of it is also found in ***kb-python***(Melsted *et al*., 2021). The purpose of the algorithm is to extract intronic regions of transcripts and write them alongside concatenated exonic regions of those transcripts. This process is used in generating RNA velocity-compatible reference sequence indices, as it allows for alignment to spliced (fully exonic transcripts) and unspliced (intronic) reference sequences. Internally, these tools use the python modules ***pyranges***(Stovner and Sætrom, 2020) and ***ngs_tools*** for genomic interval arithmetic respectively. Briefly, the algorithm parses a GTF file, extracts all exonic intervals, and computes all missing intronic regions between those exonic intervals for each transcript. ***pyroe*** also includes a step to merge intronic regions between transcripts as gene isoforms have high intronic redundancy, and so their output intronic regions are merged over those redundant intervals. Finally, these intervals are used to query an indexed fasta file and the sequences are written to a fasta file output alongside a mapping of transcript id to gene id.

The process of parsing a GTF, calculating intronic regions, and querying an indexed fasta can be represented simply using the ***bedrs*** API - resulting in a highly efficient algorithm that minimizes run-time costs and memory overhead (Figure 4). The ***bedrs*** implementation of the splici algorithm shows a run-time speedup of greater than 500x of ***kb*** and a speedup of 265x from ***pyroe***. By implementing the core functionality of genome arithmetic generically with ***bedrs***, we can separate the interval logic from the usage and development of a tool being built - and focus on building applications that are performant and expressive.

**Figure 4.**
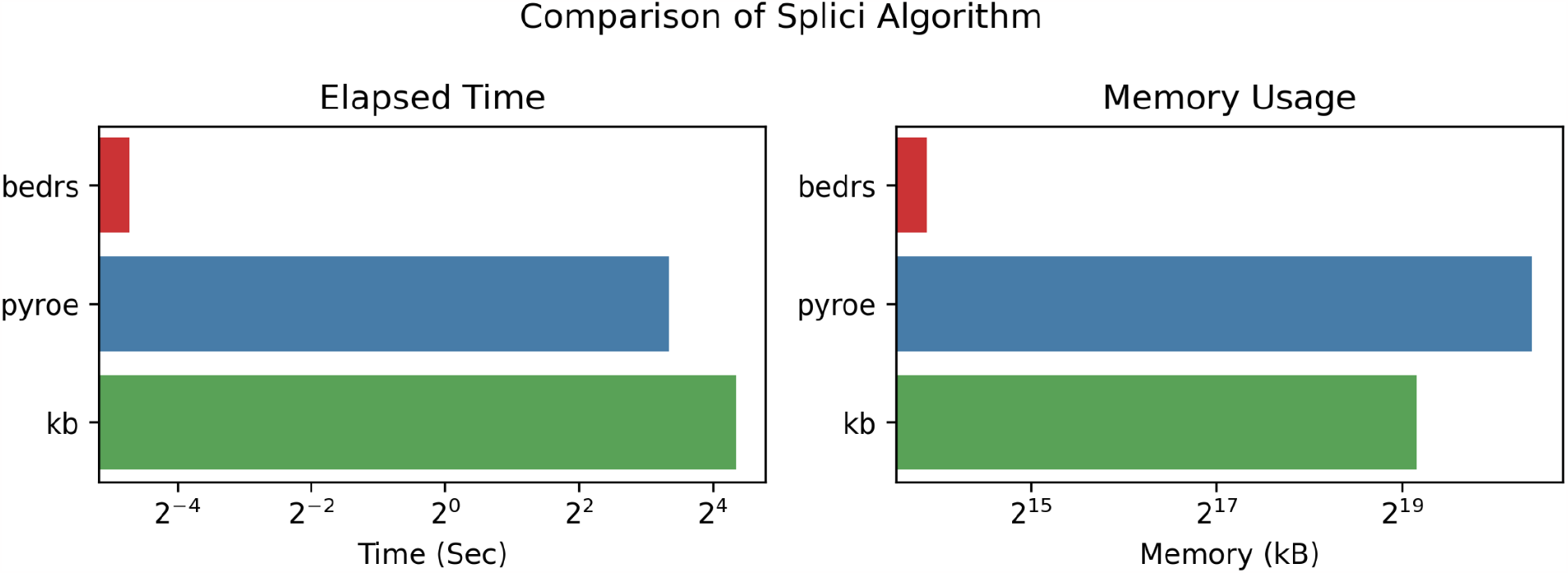
Comparison of splici algorithm runtimes written with ***bedrs*** compared to ***pyroe*** and ***kb***. Tests done using 130 random genes. On the left the elapsed time in seconds is shown. On the right the peak-memory usage for the tool is shown in kilobytes.

### Performance as a function of interval set size

The performance of ***gia*** with in-place operations on small to large interval sets is significantly higher than ***BEDTools*** and ***BEDOPS*** - but in the cases where the number of intervals is very large, the computational overhead of loading the interval set in-place becomes too large.

Where applicable, ***gia*** implements streaming operations, which keeps the memory usage flat as intervals are kept only as long as they are needed for their operation. This requires the inputs to be pre-sorted but is also more efficient when analyses require intervals to be piped between multiple operations.

We observe that ***gia*** shows lower peak memory usage than ***BEDTools*** across all interval set sizes, and shows lower peak memory usage than ***BEDOPS*** up to 10^5^ intervals (Figure 5). However, the streamed version of ***gia*** shows considerably lower peak memory usage than ***BEDTools*** and ***BEDOPS*** globally. At 10^4^ intervals, the peak memory usage is 2800 kb and 2192kb for the in-place and streamed implementations of ***gia*** respectively. At this interval size they are fairly comparable to the streamed version of ***BEDTools*** but are greater than 2x more memory efficient than the in-place version of ***BEDTools***. At this interval size as well both methods are more memory efficient than ***BEDOPS***, with a memory efficiency increase of 3.5x and 4.5x for the in-place and streamed implementations of ***gia*** respectively. The streamed implementation of ***gia*** shows a mean peak memory usage of 2223 kb, a 4.4x improvement over the peak memory usage of ***BEDOPS*** which has a consistent peak memory usage of 9887 kb.

**Figure 5.**
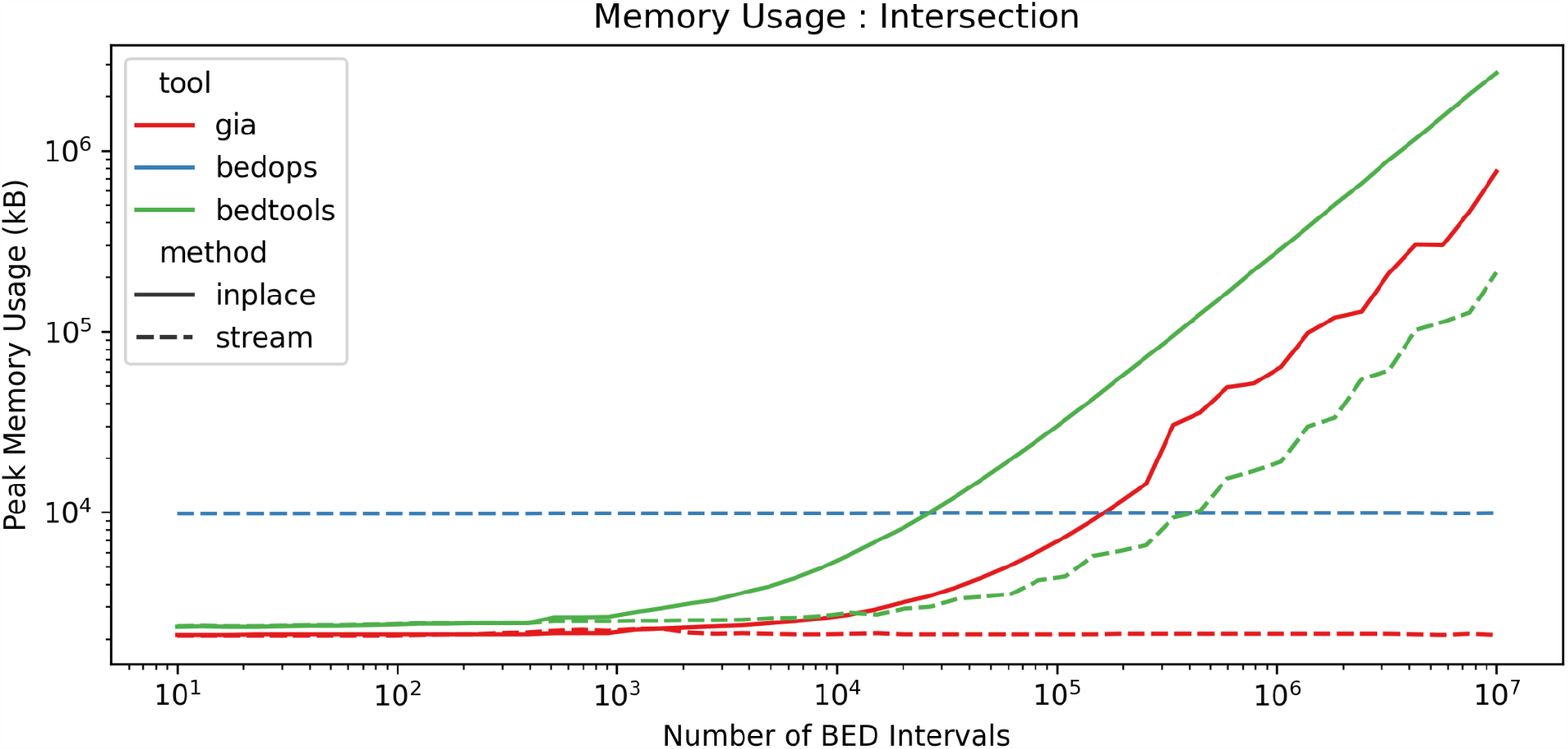
Peak-memory usage is shown for all tools implementation of intersection with in-place memory and streamable options. The number of intervals was measured over the log base 10 range of (1, 7). Memory is measured in kilobytes.

The elapsed time of the in-place implementation of ***gia*** intersection also outperforms all other methods until interval set sizes reach 4×10^5^ intervals (Figure 6). At that point the overhead of reading all the intervals and sorting becomes too great and it becomes more efficient to pre-sort the intervals and stream them. The streamable implementation of ***gia*** performs best globally and the elapsed time shows near-linear growth while in-place methods show exponential growth. ***gia*** also demonstrates an improved performance over ***BEDOPS*** in its streamed version and performs better than ***BEDOPS*** with its in-place version for even large interval set sizes.

**Figure 6.**
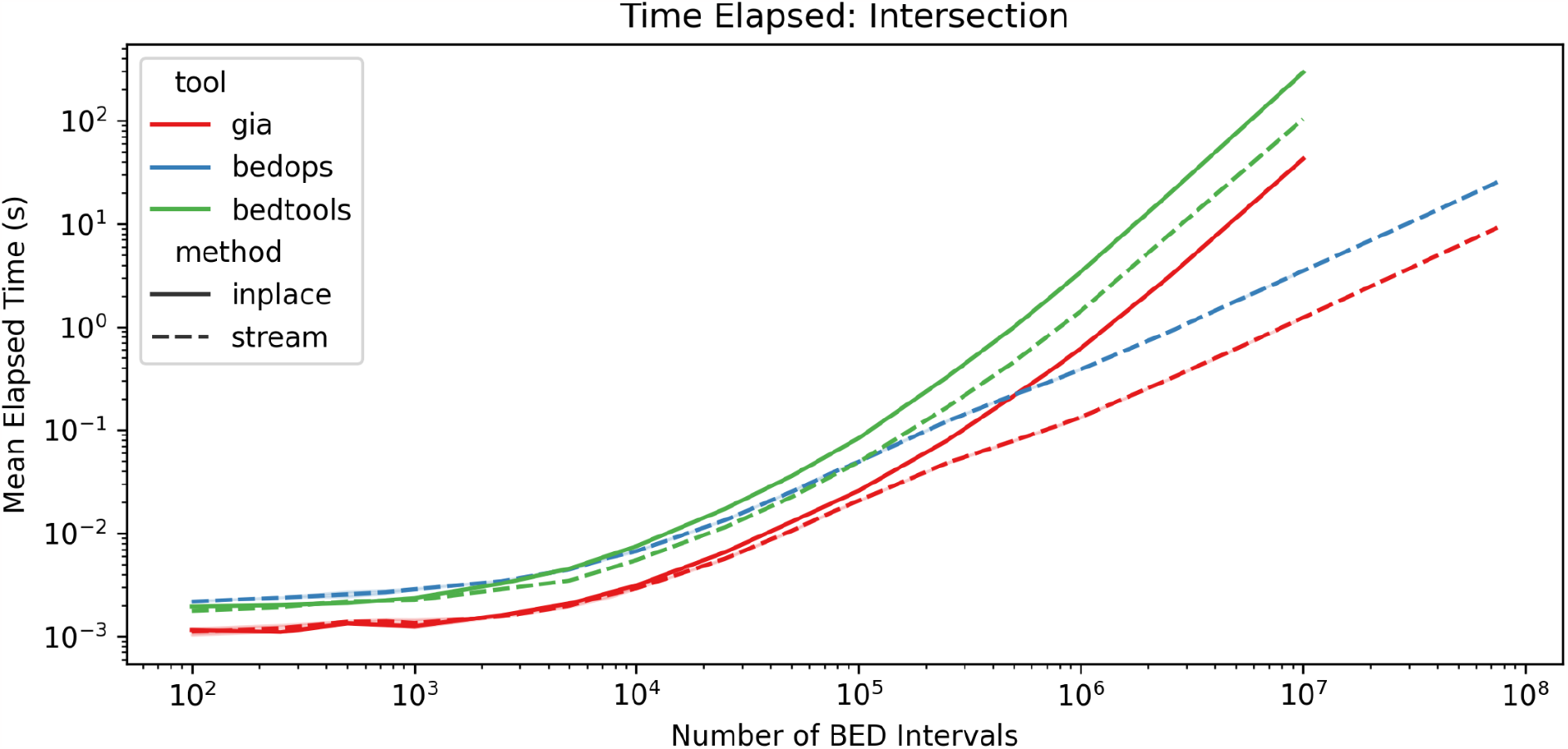
Elapsed run-time is shown for all tools implementation of intersection with in-place memory and streamable options. The number of intervals was measured over the log base 10 range of (2, 8).

## Discussion

Computing interval arithmetic is a core utility in many bioinformatics analyses and is a high-volume task in both local and remote development and infrastructure. Reducing the computational and memory overhead can drastically reduce the amount of time required for high-throughput genomic sequencing analysis, as well as reduce the cost and overhead of managing interval arithmetic requests on public and private facing servers. ***gia*** provides practical and computationally efficient methods for genomic interval arithmetic, facilitates common tasks, and can be scaled to arbitrarily sized inputs. It displays performance increases ranging from 2x to 20x for all operations compared to ***BEDOPS*** and ***BEDTools. bedrs*** provides a developer-friendly library for genome interval arithmetic and aims to open up the algorithms and data structures of genome interval arithmetic to arbitrary types and algorithms. Together they provide a proof of concept for genome interval arithmetic in rust, a practical analysis tool, and a framework for generic interval operations in bioinformatics.

### Software

***bedrs*** is free, open-source, and available at https://github.com/noamteyssier/bedrs. ***gia*** is free, open-source, and available at https://github.com/noamteyssier/gia. At time of submission the operations developed are: *closest, complement, extend, get-fasta, intersect, merge, random, sample, sort, subtract*. The ***splici*** algorithm is available as a command-line tool at https://github.com/noamteyssier/splici. The underlying indexed fasta query was developed as a modular library dependency of ‘*gia get-fasta****’*** and available at https://github.com/noamteyssier/faiquery.

## Author Contributions

N.T. designed and developed the methods, performed the analyses, and wrote the manuscript.

M.K. and H.G. provided funding and feedback. All authors read and approved the final manuscript.

## Supplementary Information

Scripts to recreate the analyses shown in this paper are accessible on github: https://github.com/noamteyssier/gia_benchmark.

## References

Danecek, P. et al. (2011) The variant call format and VCFtools. Bioinformatics, 27, 2156–2158.

Eilbeck, K. et al. (2005) The Sequence Ontology: a tool for the unification of genome annotations. Genome Biol., 6, R44.

He, D. et al. (2022) Alevin-fry unlocks rapid, accurate and memory-frugal quantification of single-cell RNA-seq data. Nat. Methods, 19, 316–322.

Kent, W.J. et al. The Human Genome Browser at UCSC.

Lawrence, M. et al. (2013) Software for Computing and Annotating Genomic Ranges. PLoS Comput. Biol., 9, e1003118.

Li, H. et al. (2009) The Sequence Alignment/Map format and SAMtools. Bioinformatics, 25, 2078–2079.

Melsted, P. et al. (2021) Modular, efficient and constant-memory single-cell RNA-seq preprocessing. Nat. Biotechnol., 39, 813–818.

Neph, S. et al. (2012) BEDOPS: high-performance genomic feature operations. Bioinformatics, 28, 1919–1920.

Peter, D. (2023) hyperfine.

Quinlan, A.R. (2014) BEDTools: The Swiss-Army Tool for Genome Feature Analysis. Curr. Protoc. Bioinforma., 47.

Quinlan, A.R. and Hall, I.M. (2010) BEDTools: a flexible suite of utilities for comparing genomic features. Bioinformatics, 26, 841–842.

Stovner, E.B. and Sætrom, P. (2020) PyRanges: efficient comparison of genomic intervals in Python. Bioinformatics, 36, 918–919.

